# Precise targeting of myotubes to muscle attachment sites patterns the musculoskeletal system

**DOI:** 10.1101/697789

**Authors:** Shuo Yang, Allison Weske, Yingqiu Du, Juliana M. Valera, Aaron N. Johnson

**Affiliations:** Department of Developmental Biology, Washington University School of Medicine in St. Louis, St. Louis, MO 63108

**Keywords:** FGF signaling, myogenesis, organogenesis, myotube guidance, Rho GTPase, actin cytoskeleton, heartless, *Drosophila* embryogenesis

## Abstract

Nascent myotubes undergo a dramatic morphological transformation during myogenesis in which the myotubes elongate over several cell diameters and choose the correct muscle attachment sites. Although this process of myotube guidance is essential to pattern the musculoskeletal system, the mechanisms that control myotube guidance remain poorly understood. Using transcriptomics, we found that components of the Fibroblast Growth Factor (FGF) signaling pathway were enriched in nascent myotubes in *Drosophila* embryos. Null mutations in the FGF receptor *heartless* (*htl*), or its ligands, caused significant myotube guidance defects. Mechanistically, paracrine FGF signals to Htl in the mesoderm regulate the activity of Rho/Rac GTPases in nascent myotubes to effect changes in the actin cytoskeleton. FGF signals are thus essential regulators of myotube guidance that act through cytoskeletal regulatory proteins to pattern the musculoskeletal system.

## Introduction

In mature skeletal muscle, myofibers are perfectly aligned with the skeleton so that muscle contractions can produce coordinated movements. During development, myotubes are directed to specific muscle attachment sites on tendons through the process of myotube guidance, and then mature into correctly aligned myofibers. Compared to our understanding of myoblast cell fate specification, migration, and fusion, relatively little is known about the molecular pathways that direct myotube guidance (Maartens and Brown, 2015).

After migrating to sites of myogenesis, myoblasts polarize and mature into nascent myotubes. Polarized nascent myotubes will extend two leading edges in opposite directions, and each leading edge navigates the extracellular environment to identify a muscle attachment site. Through this process of myotube guidance, a single myofiber will be attached to two tendons at the end of myogenesis. The intracellular pathways that reorganize the myotube cytoskeleton during guidance have been characterized in some detail. For example, the RNA binding protein Hoi polloi regulates the actin cytoskeleton by modulating Tropomyosin expression (Williams et al., 2015), and the Rho GTPase activating protein Tumbleweed, in combination with the kinesin Pavaroti, reorganizes the microtubule cytoskeleton (Guerin and Kramer, 2009). Although dynamic changes to both the actin and microtubule cytoskeletons are essential for myotube guidance, the extrinsic inputs that regulate cytoskeletal dynamics to guide myotube leading edges to the correct muscle attachment sites remain incompletely understood.

During *Drosophila* embryogenesis, Slit-Robo signaling acts as both a chemoattractant to initiate myotube elongation and as a repulsive cue to prevent myoblasts from accumulating at the ventral midline (Kramer et al., 2001). A second signaling pathway that regulates myotube guidance is directed by the orphan transmembrane receptor Kon-tiki (Kon). Kon functions through the intracellular adaptor protein Grip and, while the precise molecular function of Grip during myogenesis is still unclear, Grip may act as a scaffolding protein to cluster active Kon complexes to the myotube membrane (Schnorrer et al., 2007). Alternatively, Grip may activate intracellular signaling pathways involving small GTPases. Although the vertebrate orthologues of the Slit-Robo and Kon-Grip signaling axes have not been characterized in the context of myogenesis, Wnt11 is required to organize and orient myotubes in the trunk myotome (Gros et al., 2009). In fact, Wnt11 is the only known signaling ligand to direct myotube morphogenesis in vertebrates.

Thirty individual myotubes are specified in each segment of the *Drosophila* embryo, and each myotube acquires a highly stereotyped morphology (Bate, 1990). Disrupting the Slit-Robo or Kon-Grip signaling pathways affects only a subset of muscles in each segment (Kramer et al., 2001; Schnorrer et al., 2007), which suggests additional extrinsic inputs are required to direct myotube guidance. To identify the putative signal transduction pathways that regulate myotube guidance, we profiled the transcriptome of nascent embryonic myotubes, and found that transcripts encoding components of the Fibroblast Growth Factor (FGF) pathway were enriched in this cell population. Null mutations in the FGF receptor *heartless* (*htl*), or in the FGF ligands *pyramus* (*pyr*) and *thisbe* (*ths*), caused dramatic myotube guidance defects. *htl* mutant myotubes that expressed Htl showed largely normal muscle morphology, which argues the role of Htl is cell autonomous. Mechanistically, the Rho/Rac guanine nucleotide exchange factor *pebble* (*pbl*) and a dominant-negative form of Rac1 both suppressed the *htl* myotube guidance phenotype. Rho/Rac GTPases are well-known regulators of the actin cytoskeleton, and Htl is required to restrict Rho/Rac activity and in turn F-actin levels in nascent myotubes. This study has identified the FGF signaling as an essential component of the myotube guidance pathway that limits Rho/Rac activity to regulate cytoskeletal changes during muscle morphogenesis.

## Results

### FGF signaling components are enriched in nascent myotubes

To uncover signal transduction pathways that direct myotube guidance, we devised a Fluorescence Activated Cell Sorting and RNA deep sequencing (FACS-seq) strategy to profile the transcriptome of nascent myotubes (Fig. 1A). *rp298.GAL4* is broadly expressed in nascent myotubes, and we collected *rp298*>*GFP* embryos at 7.5-10.5hr after egg lay (Stage12-13), sorted GFP-positive myotubes and GFP-negative control cells, and isolated RNA for deep sequencing. This analysis identified 238 transcripts that were significantly enriched in nascent myotubes (Table S1), and the enriched transcripts clustered with a number of Gene Ontology (GO) terms associated with muscle development and function including *muscle cell differentiation* (GO:0042692), *myofibril* (GO:0030016), and *neuromuscular junction* (GO:0031594; Fig 1B, Table S1), suggesting our FACS-seq approach accurately identified transcripts essential to the myogenic lineage.

**Figure 1.**
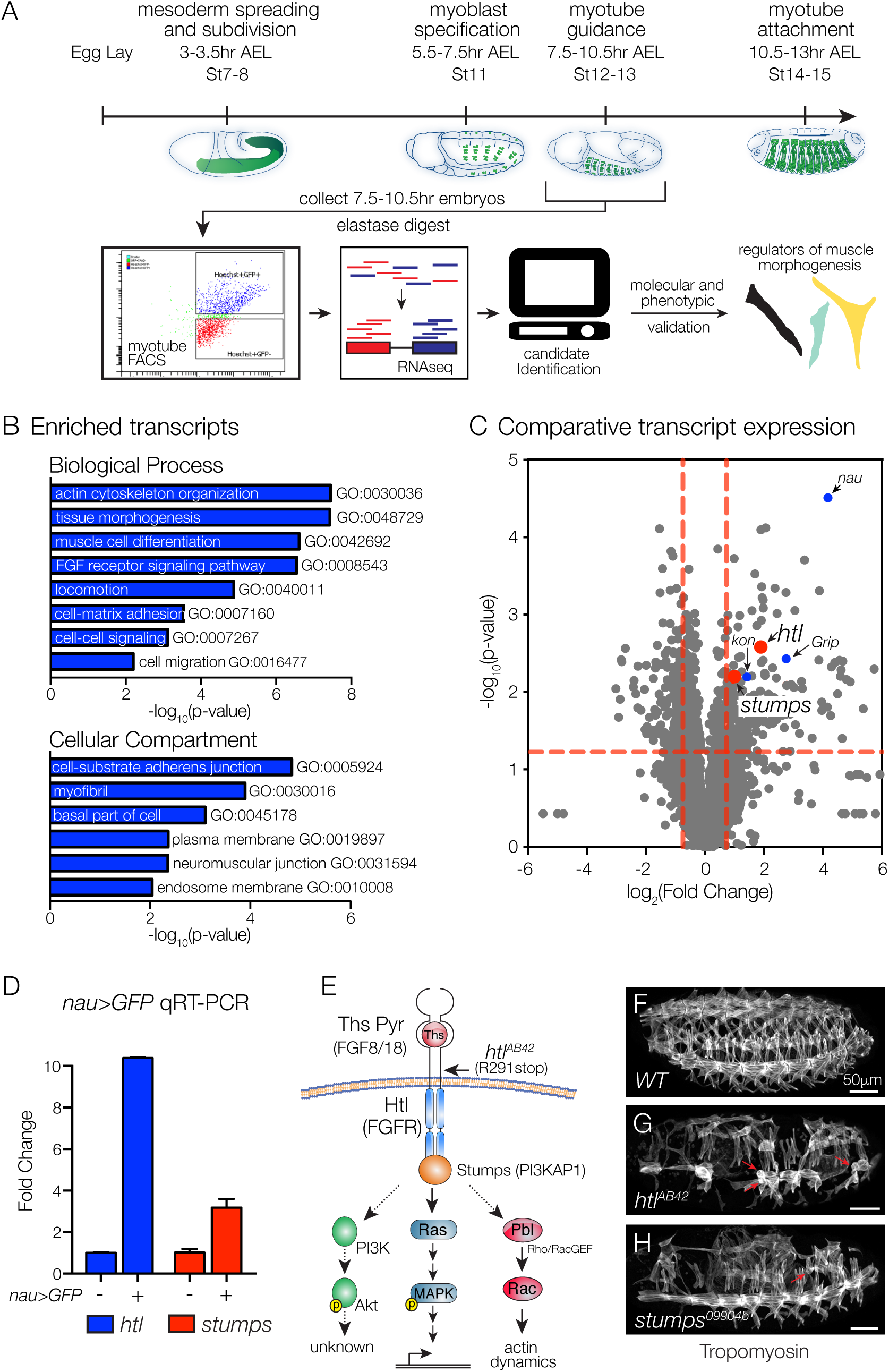
FGF signaling components are enriched in nascent myotubes. (A) Experimental design. Nascent myotubes that expressed *rp298*>*eGFP* were FACS sorted and deep sequenced (FACS-seq). Candidate genes were then tested *in vivo* to identify regulators of myotube guidance. (B) GO analysis of transcripts enriched in GFP+ myotubes compared to GFP-cells. (C) Volcano plot of transcripts identified in nascent myotubes. Each data point represents the average values for a single transcript from three biological replicates. (D) Molecular validation of FACS-seq. Nascent myotubes that expressed *nau*>*eGFP* were FACS sorted and transcript expression in sorted cells was assayed by qRT-PCR. *htl* and *stumps* transcripts were significantly enriched in GFP+ myotubes compared to GFP-cells. (E) Known FGF signaling components involving Htl. Vertebrate orthologues are given in parentheses. Indirect or putative interactions are shown with dotted lines. (F-H) St16 embryos labeled for Tropomyosin to visualize the body wall musculature. *htl*^*AB42*^ (G) and *stumps*^*09904b*^ (H) embryos showed multiple body wall muscle defects, including rounded myotubes that are indicative of myotube guidance defects (red arrows). Embryos in this and subsequent figures are oriented with anterior to the left and dorsal to the top.

One interesting cluster from the GO analysis associated with the term *FGF receptor signaling pathway* (GO:0008543; Fig 1B), and included transcripts that encode the FGF receptor Heartless (Htl), the Htl intracellular adaptor protein Stumps, and multiple enzymes that direct the synthesis of Heparin Sulfate Proteogylcans, which act as FGF co-receptors. To further analyze our FACS-seq results, we generated a scatter plot that compared the degree of statistical significance in expression with the magnitude of fold change for transcripts in the sorted and control cell populations (Fig. 1C). *kon-tiki* (*kon*) and *Grip* encode signal transducing proteins known to regulate myotube guidance (Schnorrer et al., 2007); *htl* and *stumps* showed similar values on our scatter plot as *kon* and *Grip* (Fig. 1C).

The most significantly enriched transcript from our FACS-seq experiment encodes the transcription factor Nautilus (Nau; Fig. 1C), a muscle identity gene that is expressed in a subset of nascent myotubes including Longitudinal Lateral 1 (LL1), Ventral Lateral 1 (VL1), and Ventral Oblique 5 (VO5). To validate our large-scale transcriptome profiling study, we used *nau.Gal4* to collect Stage12-13 *nau*>*GFP* embryos, and sorted GFP-positive myotubes and GFP-negative control cells. By quantitative real-time PCR (qPCR), we confirmed that *htl* and *stumps* were significantly enriched in Nau-expressing myotubes (Fig. 1D). Overall, our FACS-seq experiment highlighted a putative role for the FGF signaling pathway in regulating myotube morphogenesis.

### Htl regulates myotube guidance

Founder cells are a specialized population of myoblasts that individually give rise to a single body wall muscle. Htl is a known regulator of founder cell fate specification (Carmena et al., 1998), so we were surprised to see that *htl* transcripts were enriched in nascent myotubes several hours after founder cell specification. We hypothesized that after FGF signals specify a subset of founder cells, subsequent FGF signals direct myotube guidance. In support of this hypothesis, embryos homozygous for the amorphic allele *htl*^*AB42*^ had a number of rounded myotubes (Fig. 1E-G), a phenotype previously associated with defects in myotube elongation (Johnson et al., 2013). Embryos homozygous for a strong loss-of-function allele of *stumps* showed a similar myogenic phenotype (Fig. 1H).

To experimentally distinguish a role for Htl during myotube guidance from its known role in directing founder cell fate specification, we used transgenic lines that are expressed in discrete founder cell populations and subsequently active in nascent myotubes and mature muscles. *5053.Gal4* is expressed in VL1 founders (Fig. 2A,B), and *htl*^*AB42*^ embryos showed an equivalent number of *5053*>*GFP+* VL1 founder cells as wild type (WT) embryos (Fig. 2C,D). Although we did observe some variability in the *5053*>*GFP* expression levels in *htl*^*AB42*^ embryos, the VL1 founder cell fate was correctly specified. Consistent with the hypothesis that FGF signals direct myotube guidance, *htl*^*AB42*^ *5053*>*GFP+* VL1 muscles showed myotube elongation and muscle attachment site selection defects (Fig. 2E-G), and these guidance defects could be suppressed by expressing Htl in *htl*^*AB42*^ VL1 muscles (Fig. 2H,I). Abdominal segments 2-7 (A2-A7) show the most reproducible muscle pattern in WT embryos (Bate, 1990), so we calculated the frequency of *htl*^*AB42*^ VL1 myotube guidance defects individually in these segments. Myotube guidance defects were not confined to any subset of segments in *htl*^*AB42*^ embryos, although A2-A4 showed a slightly higher frequency of guidance defects than A5-A7 (Fig. 2J). These studies indicate that Htl acts cell autonomously to direct VL1 myotube guidance and not VL1 founder cell fate specification.

**Figure 2.**
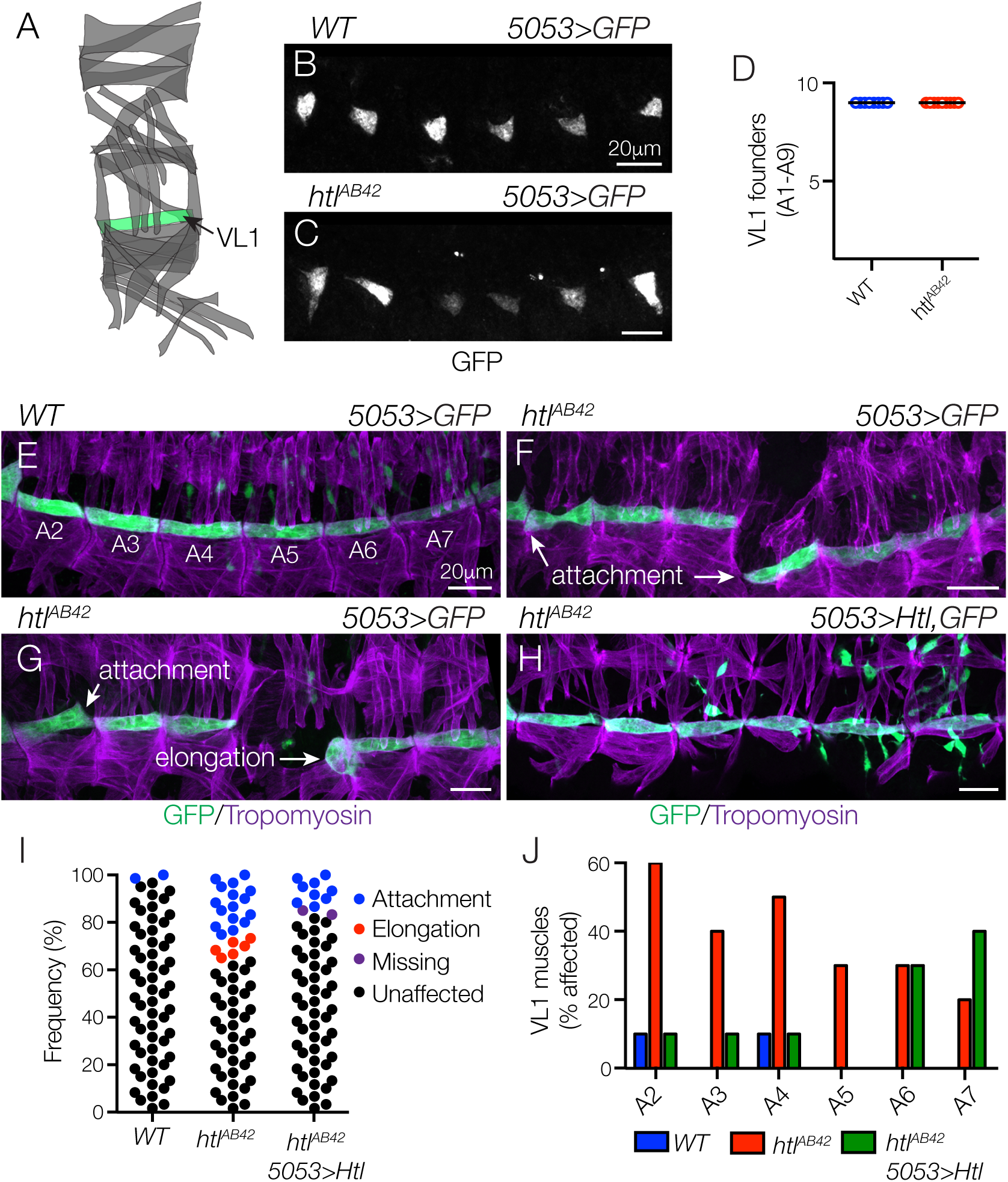
Htl acts cell autonomously to direct myotube guidance. (A) Diagram of the body wall muscles in a single embryonic segment (A2-A7). (B,C) St12 *5053*>*eGFP* embryos labeled for GFP to visualize VL1 nascent myotubes. The number of nascent myotubes specified in WT (B) and *htl*^*AB42*^ (C) embryos was equivalent (D). (E-H) St16 *5053*>*eGFP* embryos labeled for GFP (green) and Tropomyosin (violet). (E) WT VL1 muscles showed a largely invariant morphology. (F,G) *htl*^*AB42*^ VL1 muscles often attached to the wrong muscle attachment site or failed to elongate. (H) *htl*^*AB42*^ VL1 muscles that expressed Htl showed normal morphology. (I) Histogram of muscle morphology. Each data point represents a single St16 VL1 muscle in segments A2-A7. (J) Histogram of muscle defects by embryonic segment (n=60 per genotype).

### Htl directs myotube guidance in multiple cell populations

A second transgenic line, *slou.GAL4*, is expressed in Dorsal Transverse 1 (DT1), Longitudinal Oblique 1 (LO1), Ventral Transverse 1 (VT1), Ventral Acute 2 (VA2), and Ventral Acute 3 (VA3) founder cells (Fig. 3A,B). In *htl*^*AB42*^ embryos, the number of *slou*>*GFP+* cells was largely comparable to WT embryos at the onset of myotube elongation, except that the VT1 founder cells were not specified (Fig. 3C,D). At the end of myogenesis, *htl*^*AB42*^ LO1 muscles associated with incorrect muscle attachment sites and a majority of *htl*^*AB42*^ LO1 muscles acquired a lateral morphology (Fig. 3E,F,H).

**Figure 3.**
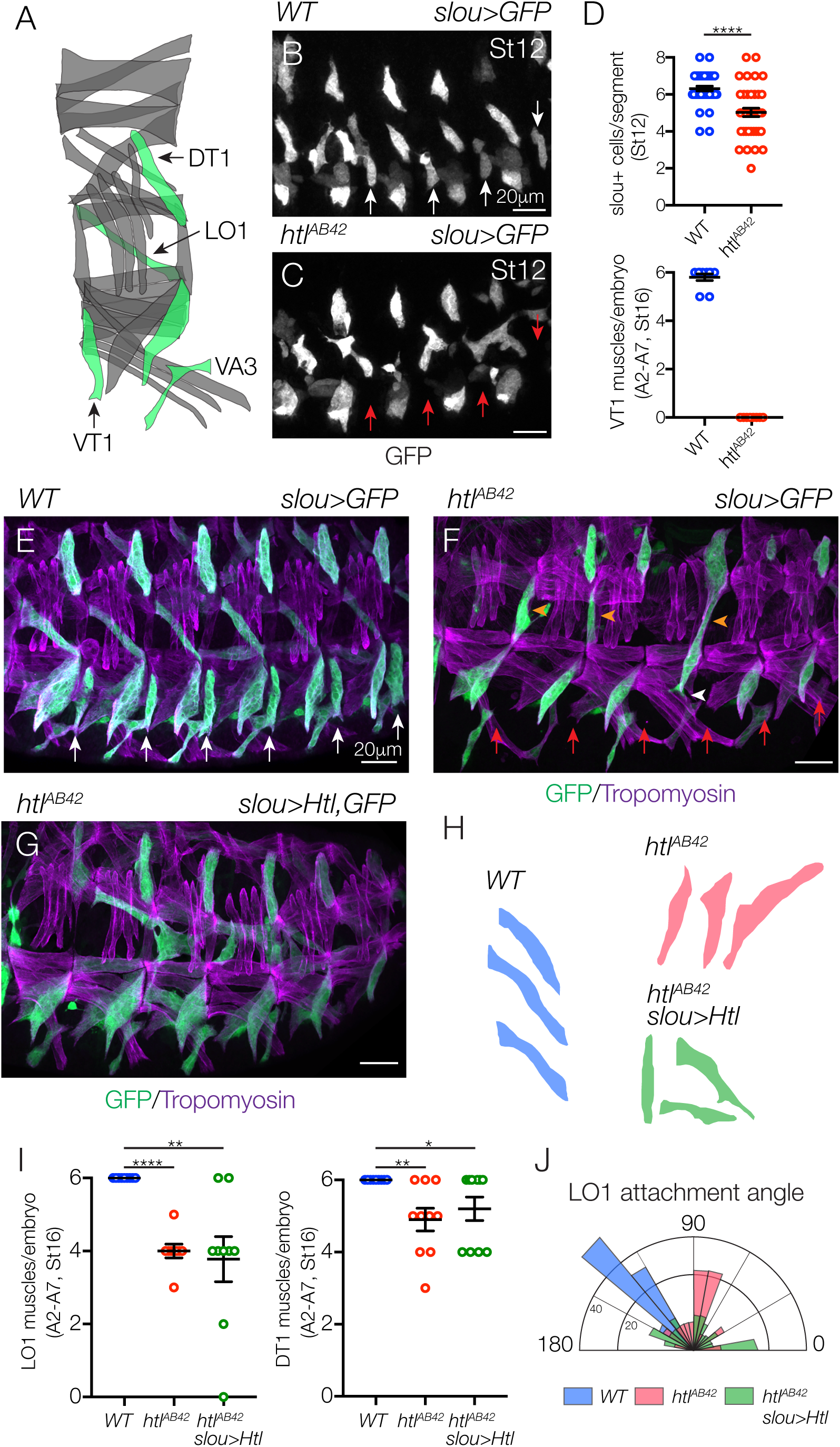
Htl acts cell autonomously and non-cell autonomously in a subset of myotubes. (A) Diagram of body wall muscles that expressed *slou.Gal4.* (B,C) St12 *slou*>*eGFP* embryos labeled for GFP. One nascent myotube per segment was absent in *htl*^*AB42*^ embryos (C, red arrows) compared to WT. (D) Muscle quantification (A2-A7). *htl*^*AB42*^ embryos failed to specify VT1 founder cells. (E-G) St16 *slou*>*eGFP* embryos labeled for GFP (green) and Tropomyosin (violet). (E) WT LO1, DT1, and VT1 muscles showed a largely invariant morphology. (F) *htl*^*AB42*^ LO1 muscles acquired a lateral morphology (orange arrowheads) and overextended ventrally (white arrowhead). *htl*^*AB42*^ DT1 muscles also showed defective morphology and VT1 muscles were absent in *htl*^*AB42*^ embryos (arrows). (G) *htl*^*AB42*^ LO1 and DT1 muscles that expressed Htl showed improved, but not WT, morphology. (H) Representative traces of LO1 muscles. (I) Number of LO1 and DT1 muscles in St16 embryos [* (p<0.05), ** (p<0.01), **** (p<0.0001); error bars represent SEM]. (J) Radial density plot of LO1 muscle attachment angles in St16 embryos. Myotube attachment angles were binned in 10° increments; each slice represents the percent of myotubes in a given bin (n≥35 muscles per genotype). See also Figure S1.

Surprisingly, *htl*^*AB42*^ embryos had significantly fewer LO1 and DT1 muscles at the end of embryogenesis than WT embryos (Fig. 3I), even though the LO1 and DT1 founder cells were correctly specified. We live imaged WT muscle morphogenesis and found that LO1 myotubes first extended a primary leading edge dorsally and perpendicular to the anterior-posterior axis. The leading edge then made a dramatic turn anteriorly and extended parallel to the anterior-posterior axis (Movie 1). This circuitous path underlies the final oblique morphology of the LO1 muscle. The secondary leading edge extended ventrally a short distance. *htl*^*AB42*^ LO1 myotubes showed three distinct guidance defects. In most cases, the primary LO1 leading edge failed to turn to the anterior, which resulted in an LO1 muscle with longitudinal morphology (Movie 1). *htl*^*AB42*^ LO1 primary leading edges also extended ventrally instead of dorsally (Movie S1), and *htl*^*AB42*^ LO1 myotubes even failed to specify a primary leading edge and both ends of the myotube extended incorrectly (Movie S2). These live imaging studies reveal that some *htl*^*AB42*^ LO1 muscles acquire unexpected morphologies and that their identification could be obscured in fixed tissues, resulting in fewer perceived *htl*^*AB42*^ LO1 muscles (Fig. 3I). In addition, *htl*^*AB42*^ LO1 leading edges often arrived at the incorrect muscle attachment site, and then moved to a different position in the embryo (Movie S1,2). This surprising result suggests that myotubes have the ability to distinguish among muscle attachment sites and that leading edge proximity to an attachment site alone is not sufficient to dictate attachment site choice.

We expressed Htl in *htl*^*AB42*^ LO1 myotubes with *slou.Gal4*, but this restricted expression was unable to completely restore *htl*^*AB42*^ LO1 muscles to WT morphology (Fig. 3G-I). To accurately quantify LO1 muscle morphogenesis, we calculated an attachment angle for individual myotubes (Fig. 3J). WT LO1 muscles had an oblique attachment angle that ranged between 120-150**°**, whereas a large proportion of *htl*^*AB42*^ LO1 muscles (45.6%) had a lateral attachment angle between 70-90**°** (Fig. 3J). Only 22.2% of *htl*^*AB42*^ LO1 muscles that expressed Htl had an attachment angle between 70-90**°** degrees (Fig. 3J), suggesting that the role of Htl during LO1 myotube guidance is partially cell autonomous.

We next considered the possibility that myotube guidance is an interdependent process and that loss of Htl activity in one myotube population might affect the morphogenesis of other myotube populations. To test this possibility, we broadly expressed Htl in all correctly specified founder cells of *htl*^*AB42*^ embryos with *rp298.Gal4*. Broad Htl expression largely restored muscle morphology in *htl*^*AB42*^ embryos, but LO1 myotubes did not acquire the stereotypically oblique morphology seen in WT embryos (Fig. S1). Since this experiment did not label individual myotubes, we could not measure specific attachment angles. These studies suggest that defects LO1 muscle morphogenesis were not rescued by Htl expression in other myotubes. More importantly, it seems the mechanisms that direct LO1 myotube guidance are more involved than those directing VL1 myotube guidance and could depend on a number of additional factors including relative levels of Htl expression or interactions with other cell types in the mesoderm (see Discussion).

### Pyr and Ths regulate myotube guidance

*Df(2R)BSC25* specifically deletes the FGF ligands *pyramus (pyr)* and *thisbe* (*ths)* (Fig. 4A), and has been used to characterize the roles of FGF signaling during mesoderm spreading and caudal visceral mesoderm (CVM) migration (Kadam et al., 2012; Stathopoulos et al., 2004). Similar to *htl*^*AB42*^ embryos (Fig. 1G), *Df(2R)BSC25* embryos (hereafter *pyr/ths* embryos) had rounded myotubes (Fig. 4B,C), and extensive defects in muscle morphogenesis (Fig. 4D). We used *slou.Gal4* and *nau.Gal4* to further compare the myogenic phenotypes between *pyr/ths* and *htl*^*AB42*^ embryos. All of the phenotypes we identified in *htl*^*AB42*^ embryos were also present in *pyr/ths* embryos, but the expressivity of the myogenic phenotypes was more severe in *pyr/ths* embryos (Fig. 4E-I, S2A-E). Interestingly, VT1 muscles were completely absent in both *htl*^*AB42*^ and *pyr/ths* embryos, while VA3 muscles were present in *htl*^*AB42*^ embryos but absent from *pyr/ths* embryos (Fig. 4E-H). These phenotypic studies argue that Pyr and Ths signal through Htl to regulate myotube guidance and to specify a subset of founder cells, but that Pyr and Ths may also signal through a Htl-independent mechanism to further direct myogenesis.

**Figure 4.**
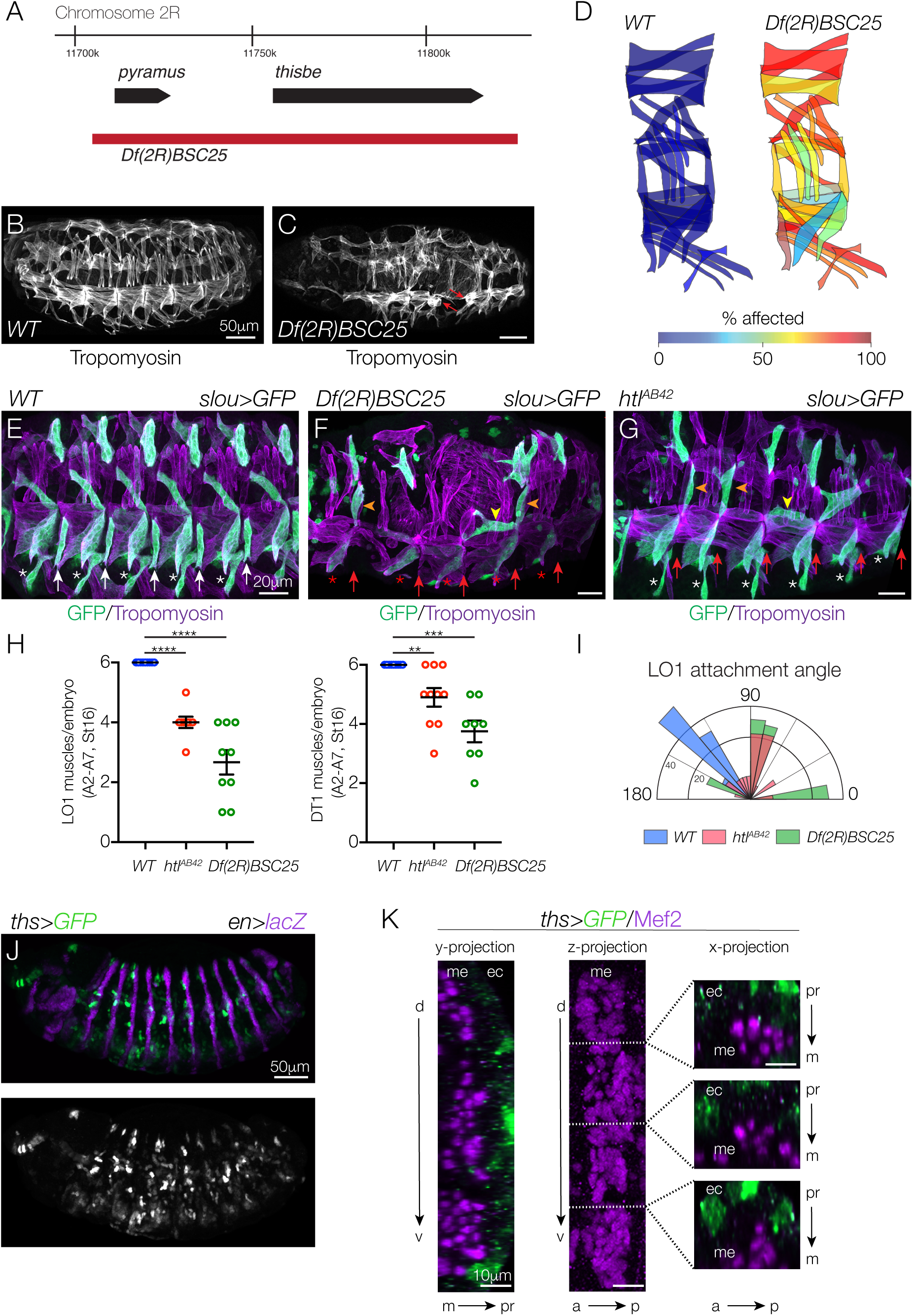
The FGF ligands Pyr and Ths direct myotube guidance. (A) Genomic organization of *pyramus (pyr)* and *thisbe (ths)*. Sequences deleted by *Df(2R)BSC25* are shown in red. (B-C) St16 embryos labeled for Tropomyosin. *Df(2R)BSC25* embryos (C) showed multiple body wall muscle defects, including missing and disorganized muscles, and rounded muscles (red arrows). (D) Heat map showing the frequency of muscle defects (n=60 segments per genotype). (E-G) St16 *slou*>*eGFP* embryos labeled for GFP (green) and Tropomyosin (violet). (F) *Df(2R)BSC25* LO1 muscles acquired lateral (orange arrowheads) and transverse morphologies (yellow arrowheads) similar to *htl*^*AB42*^ LO1 muscles (G). VT1 muscles (arrows) and VA3 muscles (asterisks) were absent in *Df(2R)BSC25* embryos. (H) Number of LO1 and DT1 muscles in St16 embryos [** (p<0.01), *** (p<0.001), **** (p<0.0001), error bars represent SEM]. (I) Radial density plot of LO1 muscle attachment angles in St16 embryos (n≥35 muscles per genotype). (J) St12 *ths*>*eGFP en.lacZ* embryo labeled for GFP (green) and lacZ (violet). *ths.Gal4* was active within and anterior to the *en* expression domain. (K) St12 *ths*>*eGFP* embryo labeled for GFP (green) and Mef2 (violet). *ths.Gal4* was active in the ectoderm (ec) but not the mesoderm (me). (a) anterior, (p) posterior, (d) dorsal, (v) ventral, (m) medial, (pr) proximal. See also Figure S2.

### Ths is expressed in the ectoderm during myotube morphogenesis

Unknown chemotactic cues from tendon cells are thought to guide myotube leading edges to muscle attachment sites (Maartens and Brown, 2015; Schnorrer et al., 2007). Ectopic expression of either Pyr or Ths at the ventral midline or in the salivary gland is sufficient to misdirect CVM migration in *pyr/ths* embryos (Kadam et al., 2012). This study suggests Pyr and Ths can be chemotactic under certain contexts. We performed a similar experiment and expressed Ths in tendon cells of *pyr/ths* embryos. However, we did not observe suppression of the *pyr/ths* myogenic phenotype or an accumulation of myotube leading edges near tendon cells (Fig. S2H,I). Importantly, tendon cells are correctly specified in *pyr/ths* embryos (Fig. S2F,G), suggesting the *pyr/ths* myogenic phenotype is not due to a lack of muscle attachment sites or overall incorrect patterning of the ectoderm. We generated *ths*>*GFP, en.lacZ* embryos and found that expression from the *ths.Gal4* knock-in (Wu et al., 2017) was not restricted to a discrete stripe in the ectoderm posterior to the Engrailed (En) expression domain (Fig. 4J), as would be expected if *ths* expression was tendon cell specific. Rather, *ths.Gal4* was expressed in limited foci in the ectoderm, both within and anterior to the En expression domain in Stage 12 embryos. We did not detect *ths.Gal4* expression in the somatic mesoderm (Fig. 4K). Although Ths expression in the ectoderm is complex, and dissimilar to that of other signaling ligands (e.g. Wingless and Hedgehog), our studies suggest that Ths expression in the ectoderm acts as a paracrine signal to regulate myotube morphogenesis.

### FGF signals are instructive

We next asked if FGF signaling could be permissive for myotubes to respond to other chemotactic cues. To test this possibility, we expressed constitutively active (CA)-Htl broadly in the founder cells of *pyr/ths* embryos, but did not observe an improvement in the *pyr/ths* myogenic phenotype (Fig. S2H,J). Thus, activating Htl alone is not sufficient to promote myotube guidance in the absence of FGF ligands.

### *htl* genetically interacts with *pbl*, a guanine nucleotide exchange factor (GEF)

FGFRs regulate multiple intracellular signaling pathways to direct cell fate specification and tissue morphogenesis, including the Mitogen-activated protein kinase (MAPK) and Protein kinase B (AKT) cascades, as well as the Rho/Rac family of small GTPases (Muha and Muller, 2013; Ornitz and Itoh, 2015). To identify the mechanism by which Htl directs myotube guidance, we first assayed phospho-MAPK (pMAPK) and pAKT levels. A well-characterized antibody has been used to identify pMAPK in *Drosophila* embryos, and although we detected pMAPK in many of the tissues previously described (Gabay et al., 1997), we did not detect appreciable pMAPK in nascent myotubes (Fig. S3A). A *Drosophila*-specific pAKT antibody has also been developed (see Materials and Methods), but this antibody does not work well for immunohistochemistry (data not shown). However, by Western Blot, we found that pAKT levels were not significantly changed in *pyr/ths* embryo lysates compared to controls (Fig. S3B). These studies suggested that FGF signaling regulates myotube guidance through MAPK- and AKT-independent mechanisms.

We next checked for genetic interactions between *htl* and the Rho/Rac GEF *pbl* (van Impel et al., 2009). Remarkably, the loss of function allele *pbl*^*3*^ dominantly suppressed overall muscle morphology defects in *htl*^*AB42*^ embryos (Fig. 5A-D), and specifically suppressed attachment site defects in *htl*^*AB42*^ VL1 muscles (Fig. 5E-H). Since *pbl* suppressed the myotube guidance phenotype in *htl* embryos, we reasoned that the role of Htl is to restrict Pbl activity in nascent myotubes. To functionally test this possibility, we expressed dominant-negative (DN) and CA-Rac1 in VL1 founder cells with *5053.Gal4*. WT VL1 myotubes that expressed DN-Rac1 showed normal morphology (Fig. 5I,M), but VL1 myotubes that expressed CA-Rac1 showed elongation and muscle attachment site defects at a frequency similar to that of *htl*^*AB42*^ VL1 muscles (Fig. 5J,M). In addition, *htl*^*AB42*^ VL1 myotubes that expressed DN-Rac1 showed fewer myotube guidance defects than *htl*^*AB42*^ VL1 muscles (Fig. 5K,M), and *htl*^*AB42*^ VL1 myotubes that expressed CA-Rac1 showed significantly more myotube guidance defects than *htl*^*AB42*^ VL1 muscles (Fig. 5L,M). In fact, *htl*^*AB42*^ VL1 myotubes that expressed CA-Rac1 most often attached to the VL3 muscle attachment sites (Fig. 5L), which we interpret as a qualitative enhancement of the *htl*^*AB42*^ phenotype. Overall, these results are consistent with a role for Htl in restricting Pbl activity to direct myotube guidance.

**Figure 5.**
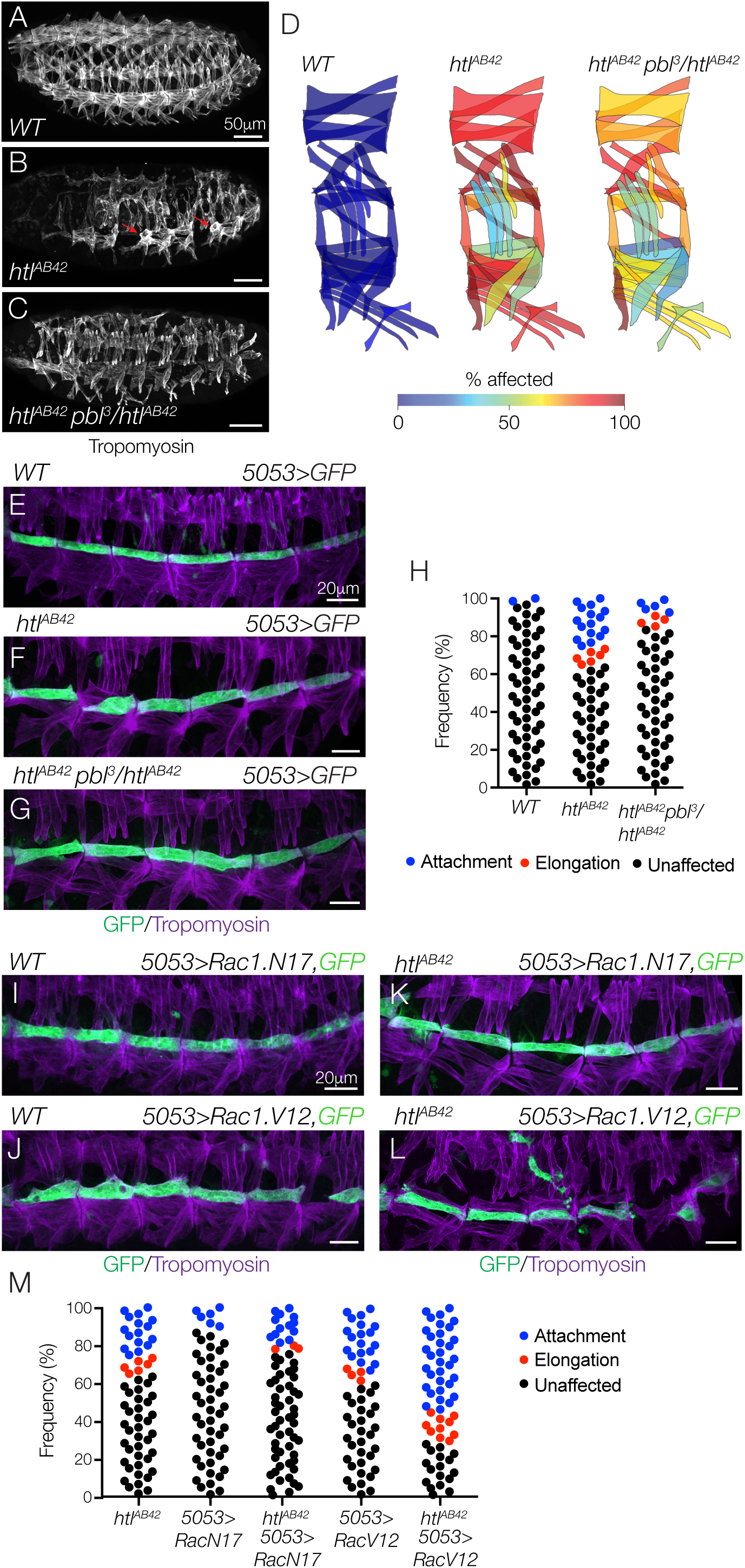
Htl regulates myotube guidance through the Pbl-Rac signaling pathway. (A-C) St16 embryos labeled for Tropomyosin. Body wall muscle defects in *htl*^*AB42*^ embryos (B, red arrows) were suppressed in *htl*^*AB42*^ embryos heterozygous for *pbl*^*3*^ (C). (D) Heat map showing the frequency of muscle defects (n=60 segments per genotype). (E-G) St16 *5053*>*eGFP* embryos labeled for GFP (green) and Tropomyosin (violet). *htl*^*AB42*^ VL1 myotube guidance defects (F) were suppressed in *htl*^*AB42*^ embryos heterozygous for *pbl*^*3*^ (G). (H) Histogram of muscle morphology. Each data point represents a single St16 VL1 muscle in segments A2-A7. (I-L) St16 *5053*>*eGFP* embryos labeled for GFP (green) and Tropomyosin (violet). VL1 myotubes that expressed dominant-negative Rac1 (Rac1.N17, I) showed normal morphology; VL1 myotubes that expressed constitutively active Rac1 (Rac1.V12, J) showed guidance defects. *htl*^*AB42*^ myotube guidance defects were suppressed in VL1 muscles that expressed Rac1.N17 (K); *htl*^*AB42*^ myotube guidance defects were dramatically enhanced in VL1 muscles that expressed Rac1.V12 (L). (M) Histogram of muscle morphology, as described in (H). See also Figure S3.

Although LO1 myotubes appeared to develop under different constraints than VL1 myotubes (Fig. 3), we hypothesized that Rho GTPases regulate a common pathway that directs both LO1 and VL1 myotube guidance. Similar to VL1 myotubes, LO1 myotubes that expressed CA-Rac1 showed severe guidance defects, whereas LO1 myotubes that expressed DN-Rac1 showed normal morphology (Fig. S3C-H). These studies suggest that Rho GTPases are essential intracellular effectors of both VL1 and LO1 myotube guidance.

### Htl limits Rho/Rac activity and F-actin assembly in nascent myotubes

To directly visualize Rho/Rac activity *in vivo*, we expressed a Rho/Rac biosensor in VL1 and LO1 myotubes. Fluorescence from the biosensor acts as a readout of Rho/Rac activity (Abreu-Blanco et al., 2014), and *htl*^*AB42*^ VL1 and LO1 myotubes showed a two to three fold increase in biosensor fluorescence compared to WT controls (Fig. 6A-E). Since the Rho/Rac family of small GTPases regulates actin dynamics (Bustelo et al., 2007), we hypothesized that F-actin levels might also be affected in *htl*^*AB42*^ myotubes. In WT VL1 and LO1 myotubes, F-actin accumulated at the leading edges and was largely excluded from the lateral membrane domains (Fig. 6F,H). In *htl*^*AB42*^ VL1 and LO1 myotubes, F-actin accumulated at the leading edges, but also along the lateral membrane domains (Fig. 6G,I). In addition, *htl*^*AB42*^ myotubes showed significantly more internal F-actin than control myotubes (Fig. 6J). From these studies we propose a model in which Htl restricts Pbl activity, Rho/Rac activation, and ultimately F-actin assembly, to direct myotube leading edges to correct muscle attachment sites (Fig. 6K).

**Figure 6.**
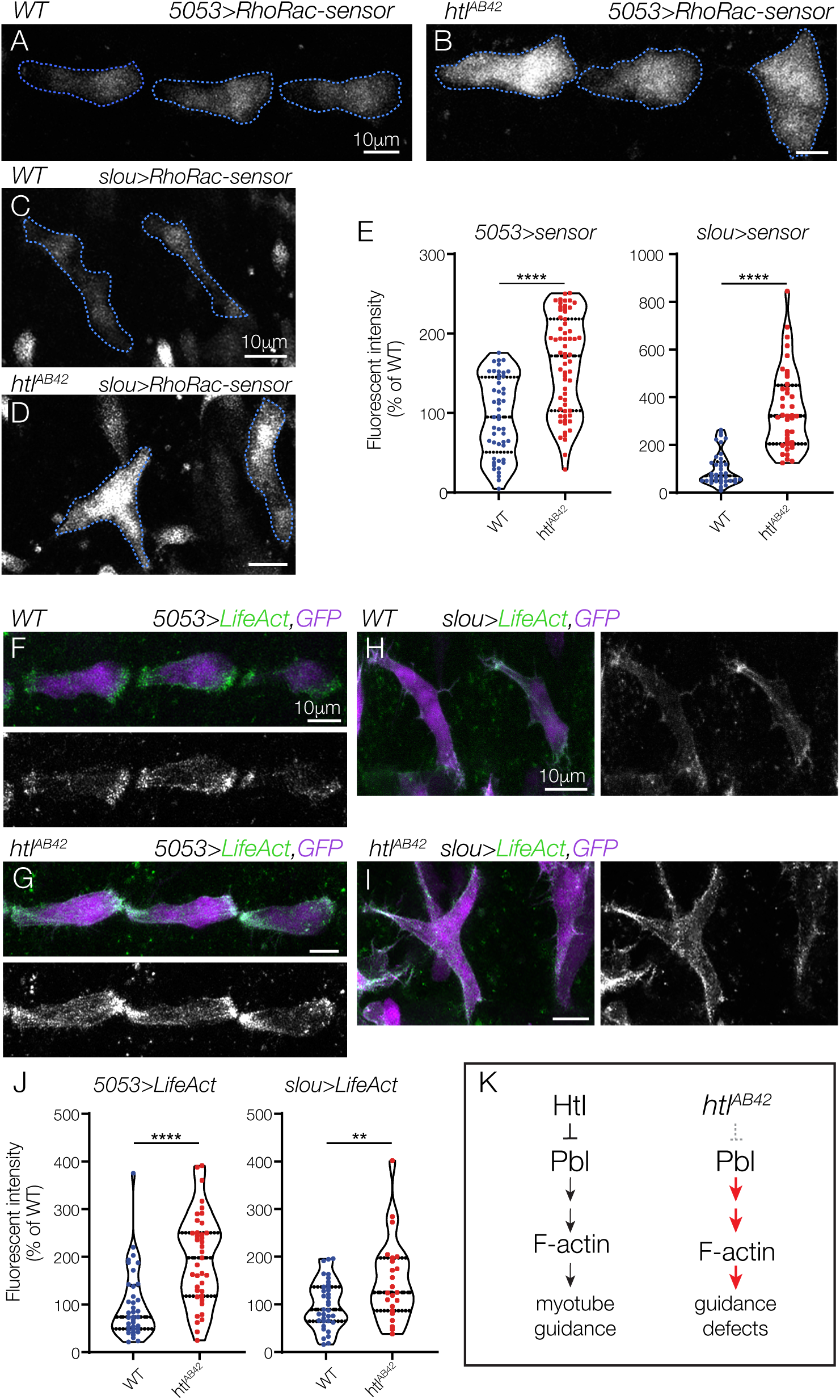
Htl restricts Rho/Rac activity during myotube guidance. (A,B) Live St13 *5053*>*RhoRac-sensor::eGFP* embryos imaged for GFP. *htl*^*AB42*^ VT1 myotubes showed more GFP fluorescence than WT VT1 myotubes. (C,D) Live St13 *slou*>*RhoRac-sensor::eGFP* embryos imaged for GFP. *htl*^*AB42*^ LO1 myotubes showed more GFP fluorescence than WT LO1 myotubes. (E) Violin plot of RhoRac-sensor fluorescence. Each data point represents a single myotube. (F,G) Live St13 *5053*>*LifeAct::RFP, eGFP* embryos imaged for GFP and RFP. F-actin accumulated primarily at the leading edges of WT VT1 myotubes (F). F-actin was not restricted to the leading edges of *htl*^*AB42*^ VT1 myotubes (G). (H,I) Live St13 *slou*>*LifeAct::RFP, eGFP* embryos imaged for GFP and RFP. WT LO1 myotubes accumulated F-actin at the leading edges (H); F-actin was not restricted to the leading edges of *htl*^*AB42*^ LO1 myotubes, and showed dramatic enrichment at the lateral membrane domains (I). (J) Violin plot of LifeAct fluorescence, as described in (E). (K) Proposed mechanism for Htl-mediated myotube guidance.

## Discussion

We have uncovered a critical role for FGF signaling in regulating myotube guidance. We found that the FGF receptor Htl acts cell autonomously in nascent myotubes to direct elongation and muscle attachment site selection. The FGF ligands Pyr and Ths are required for myotube guidance and appear to acts as paracrine signals from the ectoderm to the developing musculature. Mechanistically, Htl acts through the Rho/Rac GEF Pbl to restrict Rho/Rac activity and in turn to regulate the actin cytoskeleton. These studies provide novel insights into the mechanisms by which FGF signaling regulates the cytoskeleton to direct cellular guidance and tissue morphogenesis.

### Functional role for Htl in myotube guidance

The Htl receptor was originally identified as a regulator of mesoderm spreading in *Drosophila* (Beiman et al., 1996; Gisselbrecht et al., 1996), and later as a component of the founder cell specification gene regulatory network (Carmena et al., 1998; Michelson et al., 1998). Our study confirms the observations that Htl regulates the specification of a subset of founder cells, in particular the VT1 founders, and identifies a novel function for Htl during myotube guidance that is genetically separable from its role in founder cell specification (Fig. 2,3). Perhaps the most surprising result from our studies of Htl during myotube guidance is that *htl* myotubes could reach an incorrect muscle attachment site and then attempt to move and locate the correct attachment site (Movie S1,2). If the mechanism of myotube guidance relied on tendon precursors to establish chemoattractant gradients that myotube leading edges simply follow to the correct muscle attachment site, then the *htl* myotubes would have established a myotendinous junction with the first muscle attachment site encountered. Since *htl* myotubes recognized some muscle attachment sites as incorrect, we speculate there is a *myotendinous code* that makes some myotube-attachment site interactions permissive and others restrictive. In this model, a single myotube would have multiple permissive attachment sites, which we most convincingly observed in *htl* CA-Rac1 VL1 myotubes (Fig. 5L), and the role of Htl is to guide a myotube leading edge to the single, correct attachment site. The molecules that govern the myotendinous code are unknown, but our prediction is that permissive myotube-attachment site interactions are regulated by heterophillic interactions between cell surface proteins that are uniquely expressed in subsets of tendons and nascent myotubes.

### Htl suppresses Rho/Rac activity during myotube guidance

FGF receptors activate multiple intracellular pathways. The vertebrate MAPK, AKT, and Rho/Rac signaling pathways are activated in response to FGF signals and all three pathways influence cell migration, cell elongation, and organ morphogenesis (Benazeraf et al., 2010; Fera et al., 2004; Harding and Nechiporuk, 2012; Huebner et al., 2016; Jeong et al., 2016; Sato et al., 2011). In *Drosophila*, the intracellular pathways downstream of FGF receptors have been remarkably understudied compared to most other signal transducing receptors (Muha and Muller, 2013). We began to fill this knowledge gap by showing that the Rho/Rac GEF Pbl is an essential effector of Htl during myotube guidance. Previous genetic epistasis studies established Pbl acts downstream of Htl during mesoderm spreading (Schumacher et al., 2004), but due to the complexity of mesoderm spreading phenotypes it was unclear if Htl was an activator or repressor of Pbl activity. More recently, Rho and Rac were shown to be direct substrates of Pbl (van Impel et al., 2009), and biosensors were developed to visualize active Rho and Rac GTPases *in vivo* (Abreu-Blanco et al., 2014). Using these tools and insights, we found that the Pbl-Rho/Rac signaling axis is negatively regulated by Htl in nascent myotubes (Figs. 5A-M,6A-E), which in turn reduces overall F-actin levels and localizes F-actin to myotube leading edges (Fig. 6F-K). In addition to these mechanistic insights, we have identified the Rho/Rac biosensor a novel readout for FGF pathway activity in *Drosophila*, which to our knowledge is the only reporter of FGF receptor activity other than pMAPK.

It remains unclear how limiting Rho/Rac activity can promote myotube elongation toward the correct muscle attachment site. One possibility is that active Htl receptor complexes accumulate along the lateral myotube membranes and inhibit Rho/Rac activity everywhere except the leading edges. However, in WT myotubes Rho/Rac activity was not restricted to the leading edges (Fig. 6A). A more likely explanation is that crosstalk among the Rho family of GTPases, which includes Rho, Rac, and Cdc42, localizes the individual family members and affects overall cytoskeletal dynamics. For example, pharmalogical inhibition of Rho caused the Cdc42 expression domain to restrict and the Rac expression domain to expand in a *Drosophila* model of wound healing (Abreu-Blanco et al., 2014). In addition, dynamic but staggered accumulation of Rho and Rac to the leading edges of migratory cells is thought to drive the cytoskeletal changes that underlie the membrane expansion and retraction necessary for migration (Machacek et al., 2009). It seems plausible that Htl could inhibit Rho/Rac activity transiently to generate waves of leading edge expansion and retraction necessary for myotube guidance, and that the Rho/Rac biosensor is not sensitive to these subtle dynamics. Alternatively, Htl could restrict Rho/Rac activity in a more static fashion to promote restricted Cdc42 accumulation at the myotube leading edges.

FGF and Rho/Rac signaling also regulate cellular elongation outside of myotube guidance. For example, FGF2 promotes mammary epithelial tube elongation in organoid cultures through Rac1- and MAPK-dependent mechanisms. In this system, Rac1 inhibition caused epithelial branches to collapse after FGF2-induced elongation, but Rac1 activation alone was insufficient to induce branch elongation (Huebner et al., 2016). Unlike myotubes, mammary epithelial tubes are comprised of dozens of cells and pERK was enriched in cells at the branch tips, which suggests cells in the mammary epithelia have a differential response to exogenous FGF2. Rac1 also has a well-characterized role in regulating the actin cytoskeleton during neurite outgrowth and axon guidance (Lundquist, 2003; Luo et al., 1994), which more closely approximates myotube elongation and muscle attachment site selection as both systems involve the morphogenesis of a single cell. FGF signaling can induce neurite outgrowth (Baum et al., 2016; Saffell et al., 1997; Williams et al., 1994), and Rac1 has been implicated in regulating neuron morphogenesis downstream of FGF2 (Park et al., 2007). Our study has drawn a number of parallels between myotube guidance and axon guidance and predicts that intracellular regulation of Rho/Rac activity may be a common mechanism by which FGF signals regulate cellular elongation and guidance.

### A non-cell autonomous role for Htl

We were surprised that *htl* LO1 myotube guidance defects could not be rescued in embryos that expressed Htl under the control of *slou.Gal4*. One explanation is that *slou.Gal4* is expressed at high levels in LO1 founder cells. In fact, *slou.Gal4* activity made it the most useful reagent for live imaging studies as we could not consistently detect *5053*>*GFP* or *nau*>*GFP* founder cells in live embryos. It is possible that *slou.Gal4* activated *UAS.Htl* at levels that are much higher than endogenous *htl*, which prevented a robust rescue of *htl* LO1 myotube guidance defects.

Alternatively, Htl could have both cell autonomous and non-cell autonomous functions during LO1 myotube guidance. For example, Htl could be required to induce substrate expression for LO1 myotube guidance. The substrates for myotube guidance have not been characterized in detail, but nascent myotubes are in close proximity to the ectoderm, to fusion competent myoblasts in the somatic mesoderm, and to multiple cell types in the visceral mesoderm. Nascent LO1 myotubes are separated from the ectoderm by three Lateral Transverse myotubes (Fig. 3A), so the substrate for LO1 myotube guidance is likely expressed in mesodermal cells. In the visceral mesoderm, Caudal Visceral Mesoderm (CVM) cells migrate along the Trunk Visceral Mesoderm (TVM), and Htl performs distinct functions in each cell type. In the CVM, Htl transduces chemotactic FGF signals for directed migration, but in the TVM Htl directs integrin expression, which is the putative substrate for CVM migration (Macabenta and Stathopoulos, 2019). FGFs also regulate substrate expression in vertebrates. FGF2-induced substrate expression dramatically enhances axon regrowth across central nervous system injuries in mammals (Anderson et al., 2018). Our studies show the primary LO1 myotube leading edge travels a circuitous route (Movie 1), and one exciting possibility is that Htl-mediated FGF signals direct the expression of substrates that guide the LO1 leading edge to its muscle attachment site.

### FGFs and muscle patterning

Ectopic Ths expression in the salivary gland is sufficient to misdirect CVM migration in *pyr/ths* embryos, which suggests Ths acts as a long-range chemoattractant in this context (Kadam et al., 2012). In contrast, myotube leading edges did not preferentially extend to Ths expressing tendon cells in *pyr/ths* embryos (Fig. S2I). Since we did not detect Ths expression in tendon cells (Fig. 5J), our studies argue that ectopic Ths expression does not act as a long-range chemoattractant to induce myotube elongation across the embryonic segment. In fact, the discrete ectodermal expression pattern of Ths supports a model in which foci of FGF expression serve as ‘well-lit’ guideposts that function at short-range to direct myotube leading edges toward the correct muscle attachment sites. In either event, the mechanisms by which Pyr and Ths direct myotube guidance appear to differ from those that regulate CVM migration.

Our study has identified a novel function for the FGF pathway during myogenesis, and has established a unique experimental framework to further investigate the discrete molecular mechanisms by which FGF signaling directs cellular guidance and the physical interactions between cells during organogenesis. While FGF signaling is known to play an important role in promoting myoblast migration out of the somitic mesoderm in vertebrates, a function for FGF signaling during muscle morphogenesis has yet to be defined. Future studies of FGF signaling in both *Drosophila* and vertebrate systems will be broadly applicable toward understanding how cell shape changes are modulated by extracellular signaling pathways and may uncover much anticipated insights into how vertebrate skeletal muscles acquire spectacular shapes to complement a myriad of body plans.

## Acknowledgements

We thank Helen McNeill and Mayssa Mokalled for critical reading of the manuscript, and Richard Cripps, Angelike Stathopoulos, and the *Drosophila* community for stocks and reagents. ANJ was supported by NIH R01AR070299, the Washington University Musculoskeletal Research Center (NIH P30 AR074992), and a Boettcher Foundation Webb-Waring Biomedical Research Award.

## Author Contributions

SY, AW, YD, JMV, and ANJ designed and performed experiments. SY and ANJ prepared the manuscript.

## Experimental Procedures

### *Drosophila* genetics

The stocks used in this study include *htl*^*AB42*^, *stumps*^*09904b*^, *Df(2R)BSC25, pbl*^*3*^, *P{UAS-htl}, P{UAS-htl.λ}, P{Gal4-tey*^*5053A*^*}, P{GMR40D04-GAL4}attP2* (*slou.Gal4*), *P{GMR57C12-GAL4}attP2* (*nau.Gal4*), *Mi{Trojan-GAL4.1}ths*^*MI07139-TG4.1*^ (*ths.Gal4*), *P{UAS-Rac1.V12}, P{UAS-Rac1.N17}, P{UAS-Lifeact-RFP}, P{UAS-Pak.RBD-GFP}30, P{UAS-eGFP}* (Bloomington Stock Center), and *P{Gal4-kirre*^*rP298*^*}* (Nose et al., 1998). *Cyo, P{Gal4-Twi}, P{2X-UAS.eGFP}; Cyo, P{wg.lacZ}; TM3, P{Gal4-Twi}, P{2X-UAS.eGFP}; and TM3, P{ftz.lacZ}* balancers were used to genotype embryos.

### Fluorescence activated cell sorting and RNA sequencing (FACS-seq)

Approximately 200mg of *rp298.GAL4, UAS.eGFP* embryos were collected 7-10hr after egg lay (AEL), and dissociated as described (Bryantsev and Cripps, 2012). The cell suspension was incubated with Hoechst (1μl/ml, Invitrogen), and sorted on a FACSAria cell sorter. Minimum fluorescent intensity gates were established for GFP and Hoechst by standard methods. GFP+, Hoechst+ cells were collected for the experimental population and GFP-, Hoechst+ cells were collected for the control population. Immediately after sorting, RNA was extracted with the RNeasy Mini kit (QIAGEN). cDNA libraries were generated with the TruSeq stranded mRNA sample library kit (Illumina) and sequenced using 50bp single-reads on the Illumina HiSeq 2000 system. Three biological replicates for each cell population were sequenced in parallel and reads were screened with a custom quality control program and mapped to the *Drosophila* genome with Genomic Short-Read Nucleotide Alignment Program (GSNAP) using the Cufflinks method.

### Bioinformatic and statistical analysis

For RNA-seq, the number of fragments per kilobase of transcript per million mapped reads (FPKM) was calculated using principal-component analysis (PCA) and the relative expression for each transcript (Baird et al., 2014). Differential open reading frame (ORF) transcription between experimental and control samples was identified by calculated fold changes (FC) in FPKMs and analysis of variance (ANOVA). Transcripts with FPKM values ≥25, FC ≥1.1, and a p-value ≤0.05 in the experimental versus control populations were considered enriched and analyzed with the Database for Annotation, Visualization and Integrated Discovery (DAVID) to cluster transcripts according to Biological Process (BP) and Cellular Component (CC). Statistical analysis of embryonic phenotypes was performed with Prism 7 software, and significance was determined with the unpaired student’s t-test.

### Immunohistochemistry, imaging, and image analysis

Antibodies used include α-Mef2 (gift from R. Cripps), α-Tropomyosin (Abcam, MAC141), α-dpERK (Millipore Sigma, MAPK-YT), α-GFP (Torrey Pines Laboratories), and α-βgal (Promega). HRP-conjugated secondary antibodies in conjunction with the TSA system (Molecular Probes) were used to detect primary antibodies. Antibody staining was performed as described (Johnson et al., 2013). All images were generated with an LSM800 confocal microscope. For time-lapse imaging, dechorionated St12 embryos were mounted in halocarbon oil and scanned at 6min intervals. For other live imaging, embryos were dechorionated, mounted in PBT, and directly scanned. Control and mutant embryos were prepared and imaged in parallel where possible, and confocal imaging parameters were maintained between genotypes throughout this study. Fluorescence analysis and muscle morphology was analyzed with ImageJ software; fluorescent values were normalized to background.

### Quantitative RT-PCR

Embryonic cells were collected by FACS, and RNA was extracted with the RNeasy Mini kit (QIAGEN). cDNA was generated using Superscript IV (Life Technologies) and qPCR was performed with SYBR Select Master Mix using an ABI Prism 7000 (Life Technologies). qPCR reactions were run in triplicate and normalized to *RpL32* expression.

### Western blotting

7-10hr AEL embryos were collected, dechorionated, and suspended in lysis buffer (20 mM Tris pH 7.5, 150 mM NaCl, 1% Triton-X, and protease inhibitors). Cells were lysed with a hand-held homogenizer, and large debris was removed by 10 min centrifugation. Protein quantification of the resulting lysates was performed with Qubit Fluorometric Quantitation (Life Technologies). Western blots were performed with α-mouse-AKT (Cell Signaling Technology, 9272) and α-*Drosophila*-pAKT (Cell Signaling Technology, 4054) as described (Mokalled et al., 2010), and imaged using the ChemiDoc XRS+ system (BioRad).

