## Supplemental Figures for "Precise targeting of myotubes to muscle attachment sites patterns the musculoskeletal system" (from another version of this preprint: version 1, 2019-07-10)

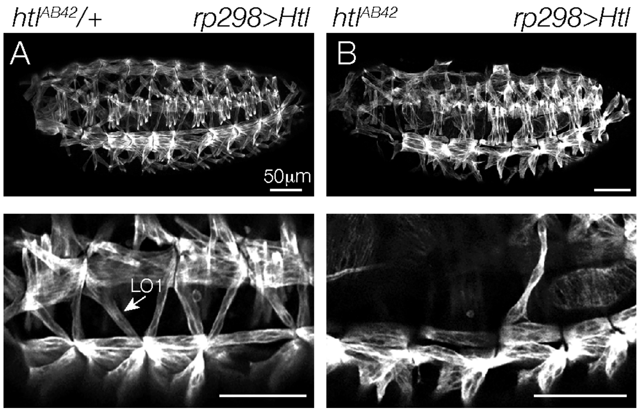


**Figure S1. Htl acts outside the body wall musculature to regulate LO1 muscle morphogenesis.** (A,B) St16 *rp298>Htl* embryos labeled for Tropomyosin. Compared to controls (A), *htl*$^{AB42}$ embryos that expressed Htl in all nascent myotubes showed LO1 muscle morphogenesis defects (B). A number of muscles in St16 *htl*$^{AB42}$ *rp298>Htl* embryos were also absent.

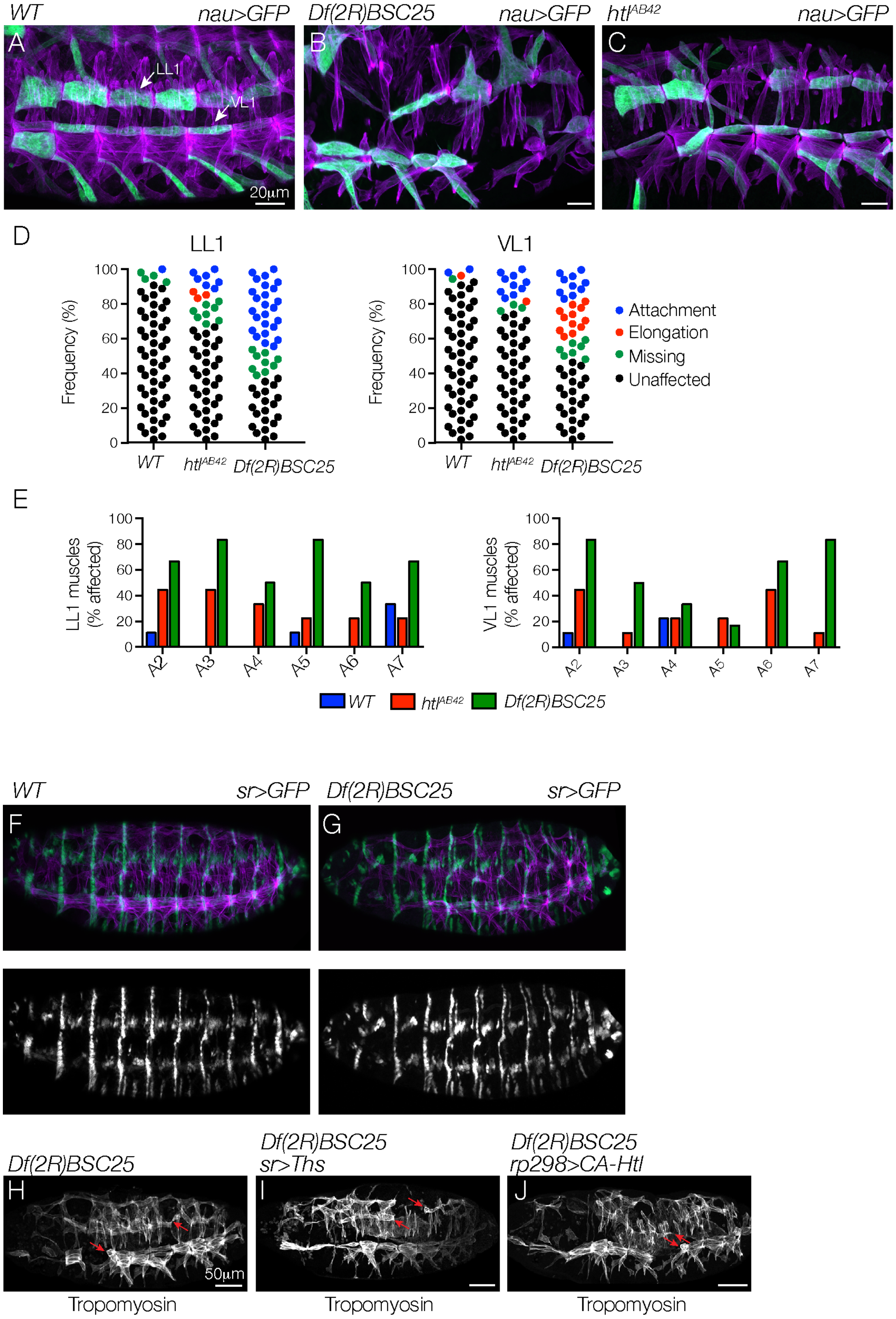
WT
nau>GFP
Df(2R)BSC25
nau>GFP
htl^AB42
nau>GFP
A
B
C
LL1
VL1
20μm
D
LL1
VL1
Frequency (%)
100
80
60
40
20
0
WT
htl^AB42
Df(2R)BSC25
Attachment
Elongation
Missing
Unaffected
E
LL1 muscles (% affected)
VL1 muscles (% affected)
A2
A3
A4
A5
A6
A7
WT
htl^AB42
Df(2R)BSC25
WT
sr>GFP
Df(2R)BSC25
sr>GFP
F
G
Df(2R)BSC25
Df(2R)BSC25 sr>Ths
Df(2R)BSC25 rp298>CA-Htl
H
I
J
50μm
Tropomyosin
Tropomyosin
Tropomyosin

**Figure S2. FGF signaling directs LL1 and VL1 myotube guidance, but not tendon cell fate specification.** (A-C) St16 *nau>eGFP* embryos labeled for GFP (green) and Tropomyosin (violet). (A) WT LL1 and VL1 muscles showed a largely invariant morphology. (B,C) LL1 and VL1 muscles in *Df(2R)BSC25* and *htl*$^{AB42}$ muscles showed attachment site and elongation defects. (D) Histogram of muscle morphology. Each data point represents a single St16 VL1 muscle in segments A2-A7. (E) Histogram of muscle defects by embryonic segment (n=54 per genotype). (F,G) St16 *sr>eGFP* embryos labeled for GFP (green) and Tropomyosin (violet). Tendon cells (marked by *sr>eGFP*) were correctly specified and maintained in *Df(2R)BSC25* embryos. (H-J) St16 *Df(2R)BSC25* embryos labeled for Tropomyosin. *Df(2R)BSC25* embryos that expressed Ths in tendon cells (I) or constitutively active Htl in nascent myotubes (J) did not show an appreciable improvement in muscle morphology compared to *Df(2R)BSC25* embryos (H).

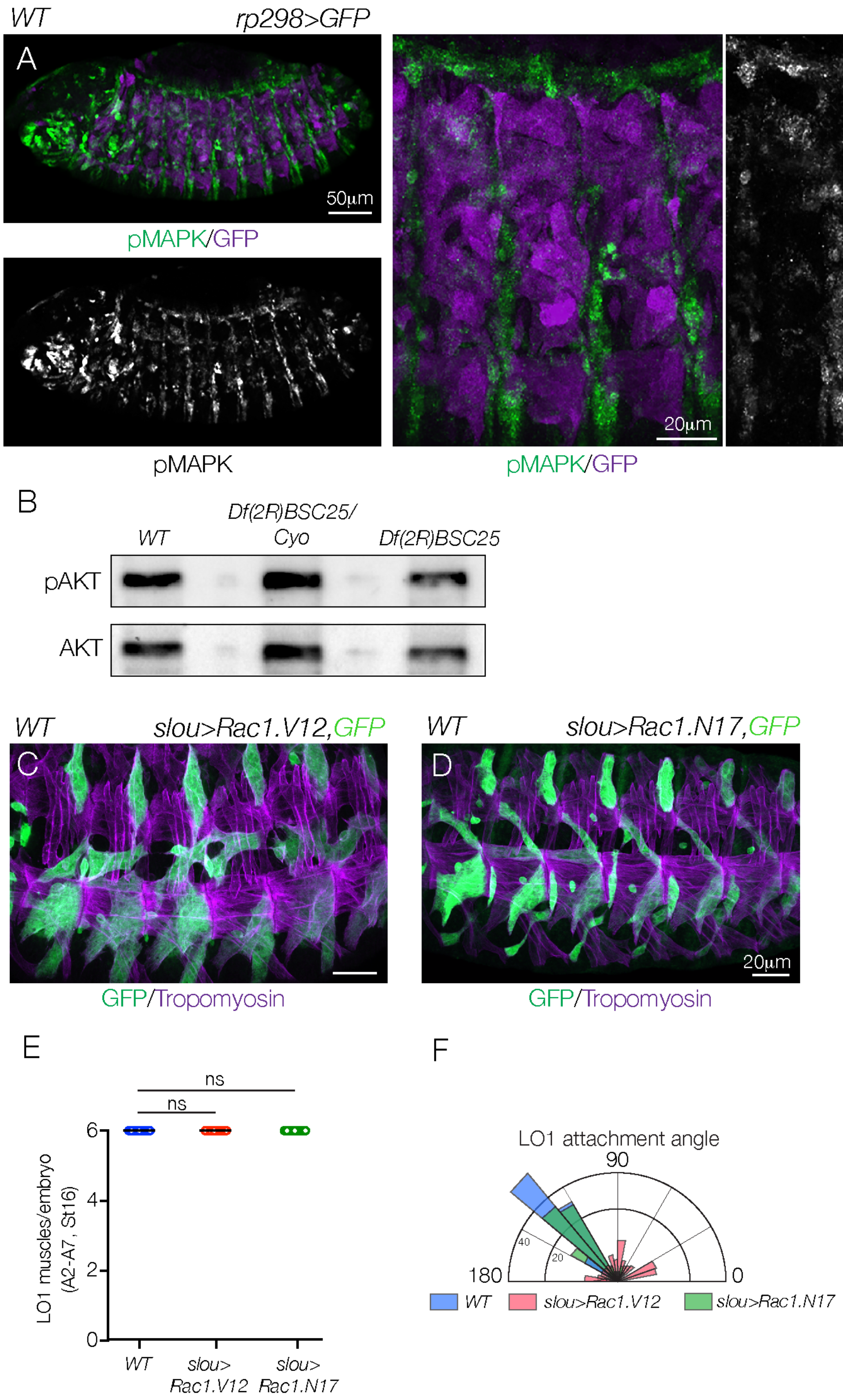


**Figure S3. Hyperactive Rac1 activity causes myotube guidance defects.** (A) St12 *rp298>eGFP* embryo labeled for pMAPK (green) and GFP (violet). Nascent myotubes did not show significant accumulation of pMAPK. (B) pAKT Western blot of St12 embryo lysates. pAKT levels were unchanged in *Df(2R)BSC25* embryo lysates compared to controls. (C,D) St16 *slou>eGFP* embryos labeled for GFP (green) and Tropomyosin (violet). LO1 myotubes that expressed constitutively active Rac1 (Rac1.V12, C) showed severe guidance defects. LO1 myotubes that expressed dominant-negative Rac1 (Rac1.N17, D) showed normal morphology. (E) Number of LO1 muscles in St16 embryos [(ns) not significant; error bars represent SEM]. (F) Radial density plots of LO1 muscle attachment angles in St16 embryos (n≥34 muscles per genotype). See Fig. 3 legend for plot details.
